# Decoding accuracies as well as ERP amplitudes do not show between-task correlations

**DOI:** 10.1101/2023.05.21.541632

**Authors:** Benedikt V. Ehinger, Hannes Bonasch

## Abstract

Some participants consistently show large, and others small activations in Electroencephalography (EEG) and other neuroimaging studies. Similarly, decoding accuracies in Brain-Computer-Interface (BCI) vary between subjects, in extreme cases labelled “BCI-Illiteracy”.

Here, we investigate whether a switch of task *within* an event-related design could be sufficient to alleviate low performance. We compare event-related-potentials (ERP) component amplitudes, as well as offline balanced decoding-accuracy based on deep convolutional networks, between seven event-related tasks. ERP effect amplitudes and decoding accuracies were correlated within all tasks, but not between any pairwise tasks. Further, 39/40 subjects had above average performance in at least one task.

Two cautious conclusions can be drawn, with the appropriate limitations of power (n=40) and the caveats of interpreting null-findings: 1) The lack of effect amplitude correlations shows that between-subject variability cannot be purely explained by a task-agnostic effects like skull thickness. 2) The lack of decoding accuracy correlations shows promise for ERP-based BCIs: replacing the task could be an effective way to combat “BCI-Illiteracy”.

## Introduction

It can often be observed that some participants show large and others small effects in EEG and other neuroimaging studies. Many explanations seem possible, e.g. in EEG there could be *task-agnostic* factors like skull thickness, known to strongly influence source localization (Antonakakis et al., 2020), varying electrode-skin impedances due to physiological different skin thereby introducing systematically more noise per subject, or idiosyncratic cortical excitability differences, influencing amplitudes of ERPs (Stephani, Hodapp, Jamshidi Idaji, Villringer, & Nikulin, 2021) -but there could as well be *task-dependent* factors, like anatomical variations resulting in detrimental dipole orientation, lacking attention or task comprehension, or task-dependent subject ability variations. The former, task-agnostic factors, should result in similar activation between tasks for any one individual, that is, task activations should show pairwise correlations.

ccA similar observation can be made not only for such encoding analyses, where brain activity is predicted by independent tasks-manipulations, but also for decoding analyses, where independent task-manipulations are predicted by brain activity^2^. Indeed, analogous to the between-subject variability in ERP amplitudes, studies of decoding models in Brain-Computer-Interfaces (BCI) show, that not every BCI can be used by any person. Around 20% of users cannot use a particular BCI at all (Allison & Neuper, 2010), often referred to as BCI-Illiteracy (Vidaurre & Blankertz, 2010; Dickhaus, Sannelli, Müller, Curio, & Blankertz, 2009) (but see (Thompson, 2019) on the term illiteracy). To date, we still do not know what factors cause between-subject variability in ERP amplitudes, nor why BCI-Illiteracy exists and how it can be alleviated.

To investigate the impact of varying task and stimulation, we estimated component amplitudes and decoding accuracies of 40 subjects, each performing seven different ERP tasks using the ERP-core dataset (Kappenman, Farrens, Zhang, Stewart, & Luck, 2020). While encoding and decoding analyses were clearly correlated within a task, we did not find any correlation between tasks. We conclude that changing the ERP task, could alleviate BCI Illiteracy. All our results are robust to within or cross-subject training, and by using robust correlations, robust to outlier subjects.

We conclude that between-subject variability cannot be explained by a task-agnostic effect, and that switching tasks within a BCI-type could provide the necessary performance boost to allow anyone to use a BCI-system.

## Methods

We used the ERP-core dataset (Kappenman et al., 2020), comprising six different ERP paradigms, which evoke seven different ERP components (N170, MMN, N2pc, N400, P300, LRP and ERN^3^). For simplicity, we refer to them as seven *tasks*. For a detailed description see the ERP-core paper (Kappenman et al., 2020).

For the **encoding analyses**, we used the preprocessed data from ERP-core, and the extraction procedure of the component peak activity as provided in https://osf.io/p3bqd/ (Kappenman et al., 2020). The only manual step we performed was the final subtraction of the per-condition component peak-estimates, resulting in a singular effect estimate per subject, per task.

For the **decoding analyses**, we used a custom pipeline (see below) starting with the raw ERP-core data, but applied similar steps as in ERP-core. We first downsampled to 250 Hz, re-referenced as described in the ERP-core paper, and finally bandpass filtered between 0.5 and 40 Hz. We only briefly mention here, that less preprocessing (only downsampling and drift-correction) as well as heavy preprocessing (additional AMICA (Palmer, Makeig, Kreutz-Delgado, & Rao, 2008), ICLabel (Pion-Tonachini, Kreutz-Delgado, & Makeig, 2019), autoreject (Jas, Engemann, Bekhti, Raimondo, & Gramfort, 2017)), to our surprise, did not show any noticeable effect on the resulting classifier accuracies. As a classifier, we used a four-layered Deep Convolutional Net as proposed by Schirrmeister et al. (Schirrmeister et al., 2017a) and implemented via Braindecode (Schirrmeister et al., 2017b). All task were 2-class problems reflecting their typical difference-waves. We used ADAM with a cross entropy loss function for optimization. We performed minimal hyperparameter tuning on the N170 task to identify useful learning rate parameters for a cosine annealing without restarts (Loshchilov & Hutter, 2017). We report balanced accuracies throughout this paper. We used stratified shuffled splits to separate training and validation data and tested both within- and across-subject training splits. While we found on average lower accuracies in the leave-one-subject-out regime, the between-task correlation patterns reported here were identical.

Finally, to compensate for potential outliers in any of the many correlation analyses, we used robust correlation coefficients using the percentage bend correlation estimator (Wilcox, 2012) as implemented in the WRS2 R package.

## Results

We analyzed encoding and decoding performance in the same subjects in seven different tasks. The encoding performance is in their task-specific typical µV-ranges (Figure 1)(Kappenman et al., 2020). We further observe typical decoding accuracies for a single subject/task between 50% and 99%, and per task averages of 60% (MMN) up to 90% (ERN). We further observed significant robust correlations between encoding and decoding analysis in all tasks with on average of 0.42 (range: 0.29 -0.58, uncorrected two-sided p-values 0.04 to *<* 0.001).

**Figure 1:**
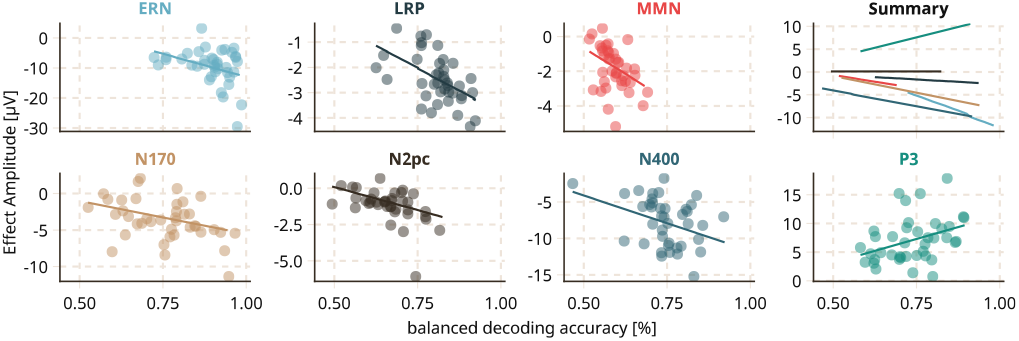
Decoding accuracies and ERP component amplitudes are correlated in all tasks.

But our main analysis regards the correlation between tasks within encoding and within decoding scores. As depicted in Figure 2, even without correcting for multiple comparisons, we only observed a single significant pairwise correlation between the N170 and the LRP, and only in the encoding data (ρ = 0.42, p=0.006, the respective decoding correlation n.s. with p = 0.051). The bulk of correlations were not significant (mean ρ:0.03 / 0.08 for encoding and decoding). This is a null-finding and has to be interpreted cautiously.

**Figure 2:**
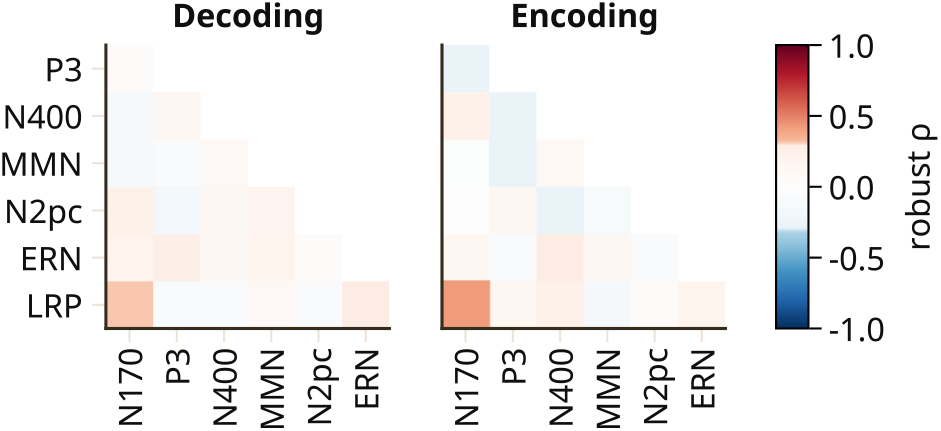
Robust correlation matrix of decoding accuracies and ERP component amplitudes. Only one correlation (LRP/N170, Encoding) is significant (no multiple comparisons correction). Compared to Figure 1, where we show within-task, between-type correlations, this figure shows the within-type, between tasks correlations.

This already hints at the fact that not any subject performs generally good or bad in all tasks. And indeed, with the exception from two subjects (Figure 3), all other subjects show above average decoding accuracy in at least two tasks. That being said, there are also *task-agnostic* effects at play, given that some subjects do show an overall better task performance (regression line in Figure 3).

**Figure 3:**
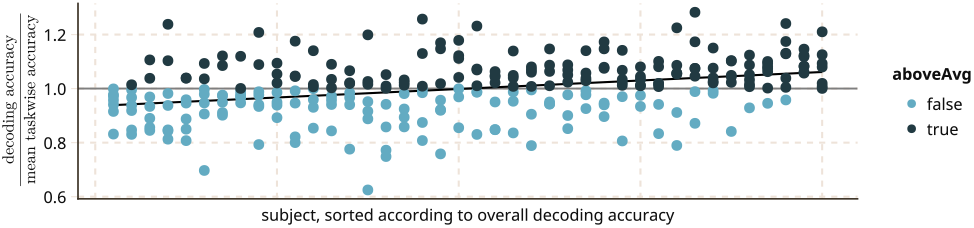
Ratio of per-subject decoding accuracy compared to task-specific mean decoding accuracy. Subjects sorted by overall mean accuracy.

## Discussion

We did not find significant correlations for neither encoding nor decoding performance across tasks.

We do have limitations on the amount of data we have, raising the question, whether our observed lack of correlation is due to sampling error in encoding/decoding performance. Overall, the influence of sampling errors should be weaker for correlations between tasks where the overall decoding accuracies are high. But this is not the case for e.g. our top-3 tasks ERN, LRP and N400. Consequently, interpreting this null-finding not as pure lack of power, but indeed as an exclusion of strong correlations, this provides at least two consequences, for 1) explaining between-subjects variability and 2) addressing BCI-Illiteracy:

1. The lack of effect amplitude correlations shows that between-subject variability cannot fully be explained by *task-agnostic* effects like skull thickness. While to some extent a *task-unspecific* effect exists, *task-specific* effects are stronger at play here. Dipole orientation is a strong contender, which would predict that lower performance should correlate with less stereotypical topographies.
2. The lack of decoding accuracy correlations shows promise for ERP-based BCIs: replacing the task could be an effective way to combat “BCI-Illiteracy”. 95% of subjects showed above average decoding performance in at least two of the seven tasks. Thus, calibrating the BCI by providing different tasks might be an effective way to make BCIs available to a broader population. One caveat needs to be highlighted here: The tasks and accuracies reported here are from standard cognitive tasks, not BCI-environments. Thus, button presses were available for decoding.

Further, this finding is interesting to the broader field of biometric identification using EEG (Sabeti, Boostani, & Moradi, 2020). In (EEG-based) biometric identification, EEG features that are unique to a subject are helpful.

Taken everything together, we see two effects at play here. A task-unspecific effect that makes some subjects high-performers and others BCI-Illiterate, but also strong task-dependent effects that might just allow everyone to use a BCI.

## Acknowledgments

Thank you to Martin Geiger for proof reading the manuscript. Decoding analyses were performed by H.B. as part of his Master Thesis. Funded by Deutsche Forschungsgemeinschaft (DFG, German Research Foundation) under Germanýs Excellence Strategy – EXC 2075 – 390740016. All necessary code for the analysis can be found under https://doi.org/10.5281/zenodo.7954914.

## Package References

**Python 3.9.16** (Van Rossum & Drake, 2009) **-PyTorch 1.13.1** (Paszke et al., 2017) **-Skorch 0.11** (Tietz, Fan, Nouri, Bossan, & skorch Developers, 2017) -**braindecode 0.7** (Schirrmeister et al., 2017b) -**pandas 1.5.3** (McKinney, 2010) – **numpy 1.24.1** (Harris et al., 2020) -**mne 1.3.0** (Gramfort et al., 2013) -**mne-bids 0.12** (Appelhoff et al., 2019) -**bids-eeg** (Pernet et al., 2019) -**autoreject 0.4.1** (Jas et al., 2017) -**scikit-learn 1.2.1** (Pedregosa et al., 2011)

**Julia 1.8.3** (Bezanson, Edelman, Karpinski, & Shah, 2017) -**AlgebraOfGraphics.jl 0.6.14** -**CairoMakie.jl 0.10.2** (Danisch & Krumbiegel, 2021) -**DataFrames.jl 1.5** -**StatsBase.jl 0.33.21** -**Pluto.jl 0.19.22 R 4.2.1** -**WRS2 1.1.4** (Mair & Wilcox, 2020)

## Footnotes

1 This paper was peer-reviewed by 11 peers and accepted for CCN23 -https://2023.ccneuro.org/. The reviewer comments were incorporated in this version

2 A simplified explanation of encoding vs. decoding models goes as follows: encoding models use *f* (*expDesign*) = *brain*, whereas decoding models use *g*(*brain*) = *expDesign. f* is typically a difference-operator or a linear regression, whereas *g* is typically some kind of 2-class classifier. Note that *g* cannot simply be inverted to receive *f* (Haufe et al., 2014)

3 LRP and ERN are extracted from the same trial at different timings.

## References

Allison, B. Z., & Neuper, C. (2010). Could Anyone Use a BCI? In D. S. Tan & A. Nijholt (Eds.), Brain-Computer Interfaces: Applying our Minds to Human-Computer Interaction (pp. 35–54). London: Springer. doi: doi:10.1007/978-1-84996-272-83

Antonakakis, M., Schrader, S., Aydin, U., Khan, A., Gross, J., Zervakis, M., … Wolters, C. H. (2020). Inter-Subject Variability of Skull Conductivity and Thickness in Calibrated Realistic Head Models. NeuroImage, 223, 117353. doi: doi:10.1016/j.neuroimage.2020.117353

Appelhoff, S., Sanderson, M., Brooks, T. L., Vliet, M. v., Quentin, R., Holdgraf, C., … Jas, M. (2019). MNE-BIDS: Organizing electrophysiological data into the BIDS format and facilitating their analysis. Journal of Open Source Software, 4(44), 1896. doi: doi:10.21105/joss.01896

Bezanson, J., Edelman, A., Karpinski, S., & Shah, V. B. (2017). Julia: A fresh approach to numerical computing. SIAM Review, 59(1), 65–98. doi: doi:10.1137/141000671

Danisch, S., & Krumbiegel, J. (2021). Makie.jl: Flexible high-performance data visualization for julia. Journal of Open Source Software, 6(65), 3349. doi: doi:10.21105/joss.03349

Dickhaus, T., Sannelli, C., Müller, K.-R., Curio, G., & Blankertz, B. (2009). Predicting BCI performance to study BCI illiteracy. BMC Neuroscience, 10(1), P84. doi: doi:10.1186/1471-2202-10-S1-P84

Gramfort, A., Luessi, M., Larson, E., Engemann, D., Strohmeier, D., Brodbeck, C., … Hämäläinen, M. (2013). MEG and EEG data analysis with MNE-Python. Frontiers in Neuroscience, 7. doi: doi:10.3389/fnins.2013.00267

Harris, C. R., Millman, K. J., van der Walt, S. J., Gommers, R., Virtanen, P., Cournapeau, D., … Oliphant, T. E. (2020). Array programming with NumPy. Nature, 585(7825), 357–362. doi: doi:10.1038/s41586-020-2649-2

Haufe, S., Meinecke, F., Gö rgen, K., Dähne, S., Haynes, J.-D., Blankertz, B., & Bießmann, F. (2014, February). On the interpretation of weight vectors of linear models in multivariate neuroimaging. NeuroImage, 87, 96–110. doi: doi:10.1016/j.neuroimage.2013.10.067

Jas, M., Engemann, D. A., Bekhti, Y., Raimondo, F., & Gramfort, A. (2017). Autoreject: Automated artifact rejection for MEG and EEG data. NeuroImage, 159, 417–429. doi: doi:10.1016/j.neuroimage.2017.06.030

Kappenman, E., Farrens, J., Zhang, W., Stewart, A. X., & Luck, S. J. (2020). ERP CORE: An Open Resource for Human Event-Related Potential Research (preprint). PsyArXiv. doi: doi:10.31234/osf.io/4azqm

Loshchilov, I., & Hutter, F. (2017). SGDR: Stochastic Gradient Descent with Warm Restarts. arXiv. (arXiv:1608.03983 [cs, math]) doi: doi:10.48550/arXiv.1608.03983

Mair, P., & Wilcox, R. (2020). Robust statistical methods in R using the WRS2 package. Behavior Research Methods, 52(2), 464–488. doi: doi:10.3758/s13428-019-01246-w

McKinney, W. (2010). Data structures for statistical computing in python. In S. van der Walt & J. Millman (Eds.), Proceedings of the 9th python in science conference (p. 51–56).

Palmer, J. A., Makeig, S., Kreutz-Delgado, K., & Rao, B. (2008). Newton method for the ICA mixture model. In Proceedings of the 33rd IEEE International Conference on Acoustics and Signal Processing (pp. 1805–1808). doi: doi:10.1109/icassp.2008.4517982

Paszke, A., Gross, S., Chintala, S., Chanan, G., Yang, E., DeVito, Z., … Lerer, A. (2017). Automatic differentiation in PyTorch. In Nips autodiff workshop.

Pedregosa, F., Varoquaux, G., Gramfort, A., Michel, V., Thirion, B., Grisel, O., … Duchesnay, E. (2011). Scikitlearn: Machine Learning in Python. Journal of Machine Learning Research, 12, 2825–2830.

Pernet, C. R., Appelhoff, S., Gorgolewski, K. J., Flandin, G., Phillips, C., Delorme, A., & Oostenveld, R. (2019). EEG-BIDS, an extension to the brain imaging data structure for electroencephalography. Scientific Data, 6(1), 103. doi: doi:10.1038/s41597-019-0104-8

Pion-Tonachini, L., Kreutz-Delgado, K., & Makeig, S. (2019). ICLabel: An automated electroencephalographic independent component classifier, dataset, and website. NeuroImage, 198, 181–197. doi: doi:10.1016/j.neuroimage.2019.05.026

Sabeti, M., Boostani, R., & Moradi, E. (2020). Event related potential (ERP) as a reliable biometric indicator: A comparative approach. Array, 6, 100026. doi: doi:10.1016/j.array.2020.100026

Schirrmeister, R. T., Springenberg, J. T., Fiederer, L. D. J., Glasstetter, M., Eggensperger, K., Tangermann, M., … Ball, T. (2017a). Deep learning with convolutional neural networks for EEG decoding and visualization. Human Brain Mapping, 38(11), 5391–5420. (arXiv:1703.05051 [cs]) doi: doi:10.1002/hbm.23730

Schirrmeister, R. T., Springenberg, J. T., Fiederer, L. D. J., Glasstetter, M., Eggensperger, K., Tangermann, M., … Ball, T. (2017b). Deep learning with convolutional neural networks for eeg decoding and visualization. Human Brain Mapping. doi: doi:10.1002/hbm.23730

Stephani, T., Hodapp, A., Jamshidi Idaji, M., Villringer, A., & Nikulin, V. V. (2021). Neural excitability and sensory input determine intensity perception with opposing directions in initial cortical responses. eLife, 10, e67838. doi: doi:10.7554/eLife.67838

Thompson, M. C. (2019). Critiquing the Concept of BCI Illiteracy. Science and Engineering Ethics, 25(4), 1217–1233. doi: doi:10.1007/s11948-018-0061-1

Tietz, M., Fan, T. J., Nouri, D., Bossan, B., & skorch Developers. (2017, July). skorch: A scikit-learn compatible neural network library that wraps pytorch [Computer software manual].

Van Rossum, G., & Drake, F. L. (2009). Python 3 reference manual. Scotts Valley, CA: CreateSpace.

Vidaurre, C., & Blankertz, B. (2010). Towards a Cure for BCI Illiteracy. Brain Topography, 23(2), 194–198. doi: doi:10.1007/s10548-009-0121-6

Wilcox, R. R. (2012). Introduction to robust estimation and hypothesis testing. doi: doi:10.2307/2669876

